# Glioma Through the Looking GLASS: the <u>G</u>lioma <u>L</u>ongitudinal <u>A</u>naly<u>s</u>i<u>s</u> consortium, molecular evolution of diffuse gliomas

**DOI:** 10.1101/196139

**Authors:** The GLASS consortium, Kenneth Aldape, Samirkumar B Amin, David M Ashley, Jill S Barnholtz-Sloan, Amanda J Bates, Rameen Beroukhim, Christoph Bock, Daniel J Brat, Elizabeth B Claus, Joseph F Costello, John F de Groot, Gaetano Finocchiaro, Pim J French, Hui K Gan, Brent Griffith, Christel C Herold-Mende, Craig Horbinski, Antonio Iavarone, Steven N Kalkanis, Konstantina Karabatsou, Hoon Kim, Mathilde CM Kouwenhoven, Kerrie L McDonald, Hrvoje Miletic, Do-Hyun Nam, Ho Keung Ng, Simone P Niclou, Houtan Noushmehr, D Ryan Ormond, Laila M Poisson, Guido Reifenberger, Federico Roncaroli, Jason K Sa, Peter AE Sillevis Smitt, Marion Smits, Camila F Souza, Ghazaleh Tabatabai, Erwin G Van Meir, Roel GW Verhaak, Colin Watts, Pieter Wesseling, Adelheid Woehrer, WK Alfred Yung, Christine Jungk, Ann-Christin Hau, Eric van Dyck, Bart A Westerman, Julia Yin, Olajide Abiola, Mustafa Khasraw, Erik P Sulman, Andrea M Muscat

## Abstract

Adult diffuse glioma are a diverse group of intracranial neoplasms associated with a disproportional large number of productive life years lost, thus imposing a highly emotional and significant financial burden on society. Patient death is the result of an aggressive course of disease following diagnosis. The Cancer Genome Atlas and similar projects have provided a comprehensive understanding of the somatic alterations and molecular subtypes of glioma at diagnosis. However, gliomas undergo significant molecular evolution during the malignant transformation. We review current knowledge on genomic, epigenomic and transcriptomic abnormalities before and after disease recurrence. We outline an effort to systemically catalogue the longitudinal changes in gliomas, the <u>G</u>lioma <u>L</u>ongitudinal <u>A</u>naly<u>s</u>i<u>s</u> Consortium. The GLASS initiative will provide essential insights into the evolution of glioma towards a lethal phenotype with the potential to reveal targetable vulnerabilities, and ultimately, improved outcomes for a patient population in need.

## 1. Introduction

Diffuse gliomas are the most frequent malignant primary brain tumors in adults.^1^ Almost all relapse despite intense treatment with surgery, radiation, and chemotherapy. The most common and most aggressive gliomas, glioblastoma (GBM), are IDH-wildtype and classified as World Health Organization (WHO) grade IV. They are characterized by a median overall survival that has remained static at around 15 months for decades even in selected clinical trial populations.^2-5^ Patients with lower-grade (WHO grades II and III) IDH mutated gliomas have a more favorable prognosis but their tumors are also lethal, as they too recur and become resistant to therapy.^1^ The standard of care for infiltrative/diffuse gliomas is maximal safe resection, followed by chemoradiation.^6^ Patients are then monitored for disease progression by imaging at regular intervals following surgery. Evaluation of disease progression is commonly guided by specific criteria (e.g. Response Assessment in Neuro-Oncology (RANO)),^7^ which rely on visual evaluation of contrast enhancement and the non-enhancing hyperintense area on T2-weighted imaging. Radiologic features sometimes do not distinguish between true tumor progression and its imaging mimicker, pseudo-progression, and disease progression cannot be reliably established based on imaging alone. Inaccurate assessment can result in premature withdrawal from a specific treatment or to continue an ineffective therapy. A further challenge in particular in the monitoring of lower-grade gliomas is the prediction of malignant transformation without another surgical intervention leading to a new histological analysis. At present, this is based on clinical progression and rather imprecise imaging signs such as contrast enhancement in a previously non-enhancing lesion or increase in size of measurable abnormalities, that are better indicators of existing imaging features rather than impending malignant transformation.

Molecular characterization of gliomas in recent years has advanced our understanding of their genesis and has identified abnormalities that allow a better classification and may be therapeutically targetable^8^. Through efforts by the Cancer Genome Atlas (TCGA),^9-14^ the genomes of over 1,100 grades II–IV gliomas have been characterized in detail, with other groups contributing further important results and datasets.^15-20^ This wealth of information has provided a detailed molecular portrait of primary glioma. The atlas, by design, focused on untreated tumors. The next frontier in glioma genomics is to understand recurrent disease. This is important, as patients generally die from tumor regrowth after therapy that becomes increasingly more resistant. Recent reports on pilot sets of paired tumors obtained before and after therapy show that there are many differences between the primary neoplasm at diagnosis and the recurrent tumor.^21^ Malignant progression of gliomas, similar to other cancers, is the result of an evolutionary process that involves reiterative cycles of clonal expansion, genetic diversification, and clonal selection under microenvironmental pressures, including overcoming antitumor immune responses.^22^ The presence of multiple cell populations with an array of different mutations is at least partly responsible for rapid induction of intrinsic resistance to therapy in gliomas.^23^ Adaptive epigenetic and phenotypic responses are equally important. The emerging understanding of this dynamic evolution of the glioma genome has major implications for cancer biology research and potential development of effective therapy. However, like the molecular landscape of primary tumors, a full understanding of the dynamic evolution of gliomas there can only be achieved through (a) profiling of sufficiently large patient cohorts to achieve statistical significance and (b) standardization across biospecimen processing and data platforms. The <u>G</u>lioma <u>L</u>ongitudinal <u>A</u>naly<u>S</u>i<u>S</u> (GLASS) consortium has been initiated to generate a comprehensive molecular and radiological portrait from paired primary and recurrent adult gliomas, including clinical patient data, to enable discovery of vulnerabilities that render the tumor sensitive to therapeutic intervention (Figure 1).

**Figure 1.**
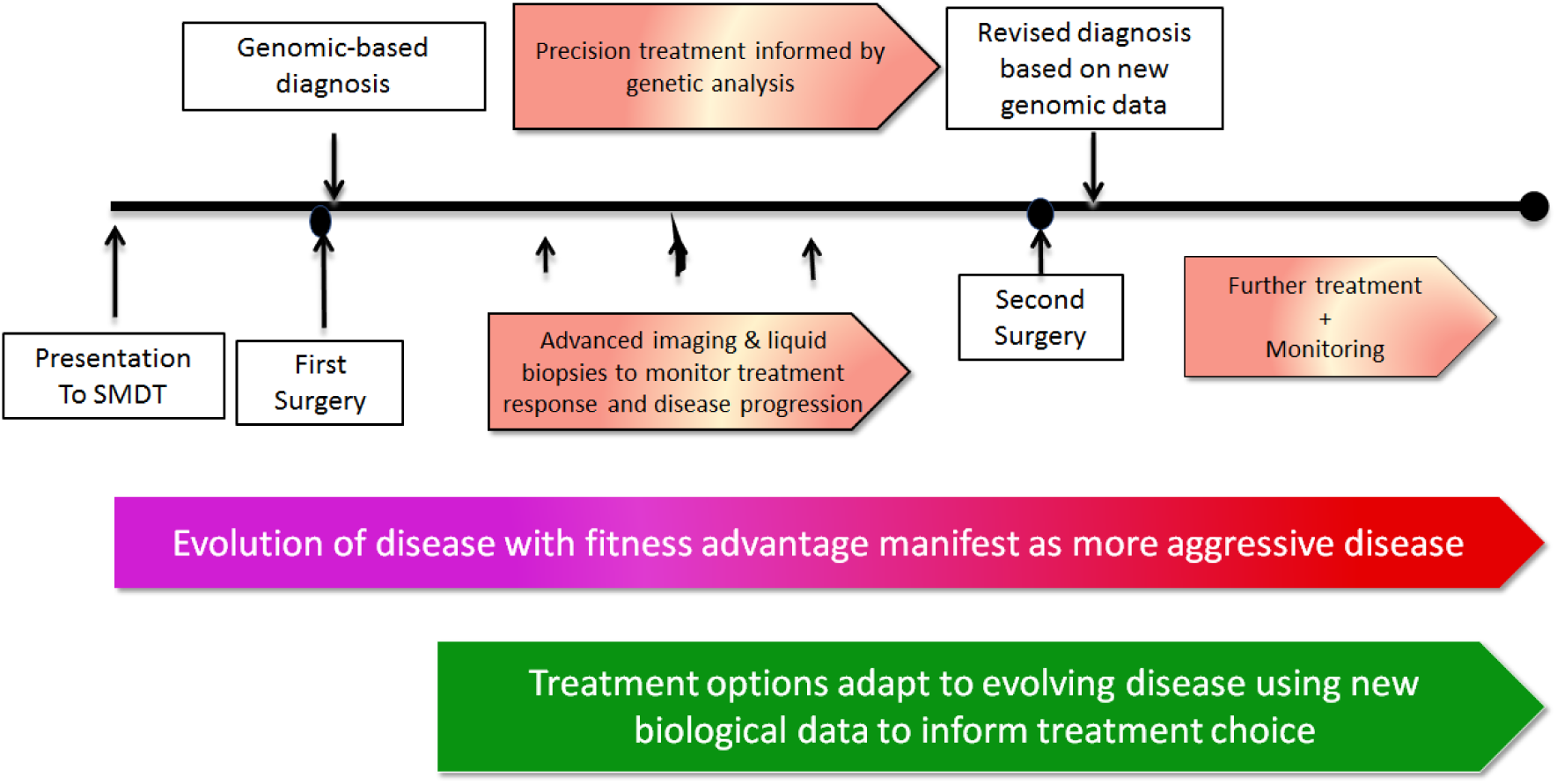
Usual course of glioma management. GLASS would improve the assessment of gliomas, particularly prediction of malignant transformation, treatment monitoring, and assessment of tumor alterations non-invasively with imaging and/or liquid biopsies. SMDT: Specialist multidisciplinary team; CSF: Cerebrospinal fluid; RT: Radiotherapy; TMZ: Alkylating antineoplastic agent Temozolomide; 1p19q: Short arm of chromosome 1 (1p) and long arm of chromosome 19 (19q); IDH: Isocitrate dehydrogenase; MGMT: O-6-methylguanine-DNA methyltransferase; TERT: Telomerase reverse transcriptase.

## 2. Molecular profiling offers new possibilities for diagnosis and therapy of gliomas

### 2.1 Clinical classification of adult diffuse glioma

For over a century, microscopic evaluation provided the gold standard for diagnosis of diffuse gliomas, and assessment of prognosis and therapeutic management of patients were based on histopathologic diagnosis.^24^ Over the past two decades, it has become clear that combination of histopathology with specific molecular characteristics of gliomas provides a more robust and objective basis for clinical stratification.^8,11,13,17,19,25-30^ Three large clinically relevant subgroups of diffuse gliomas in adults can be defined on the basis of firstly, presence or absence of mutations in the isocitrate dehydrogenase (*IDH*) 1 gene or *IDH2* gene and secondly, within the category of IDH mutated glioma, either complete 1p/19q codeletion (combined loss of the short arm of chromosome 1 and the long arm of chromosome19) or *TP53* mutation and *ATRX* loss.^11,13,17,19,25,27-30^ Most WHO grade II and III diffuse astrocytomas and oligodendrogliomas are IDH mutant, and about a third of these contain 1p/19q codeletion. In contrast, approximately 95% of glioblastomas are IDH-wildtype.^14^ IDH mutations are centered on codon 132 of *IDH1* and 172 of *IDH2*. About 90% of IDH-mutant gliomas are R132H *IDH1* mutant.^31,32^ The protein resulting from this mutation can be recognized by a highly specific and sensitive antibody available for clinical practice^31^, or through targeted sequencing. Remaining IDH-mutant cases may carry alternate R132 alleles such as R132C or variants in *IDH2*. Expert consensus on how these molecular data should be implemented in routine clinical practice^26^ led to revision in 2016 of the WHO 2007 classification of CNS tumors.^8^ These revisions indeed integrate histopathological and molecular data 8 and for the first time, this scheme provides integrated data for diagnosis, prognostic grading, and guidance of therapeutic decisions.^33,34^ In the revised classification, mutations in genes such as *TP53*, and *ATRX* can also be added to support or refine the diagnosis but are not mandatory. However, even this greatly improved classification system is predicated on primary, untreated disease; it is still unclear how these molecular markers impact the biology and prognosis of post-therapy, recurrent glioma. The promoter DNA methylation status of the gene *MGMT* is predictive of response to temozolomide therapy in primary tumors and this status is thought to be mostly stable between primary and recurrent disease ^35^. The value of re-testing MGMT status after disease progression is debatable and a methylated MGMT promoter continues to predict treatment response at this stage.

### 2.2 Intratumoral heterogeneity in primary gliomas

Cancer results from a single normal cell that has acquired molecular alterations providing it with a proliferative advantage. In glioma, the most frequent somatic abnormalities are thought to be founding events. This included the mutations in the IDH genes and mutations in the promoter of telomerase reverse transcriptase (*TERT*) gene, especially characteristic of IDH-wildtype GBM and IDH-mutant, 1p/19q-codeleted gliomas.^25^ Major aneuploidy, such as the 1p/19q co-deletion but also whole chromosome 7 gain and chromosome 10 loss which is common in IDH-wildtype glioma, are also thought to be glioma originating alterations. ^36-38^ The divergence in somatic abnormality profiles between the three major glioma subtypes, IDH-wildtype, IDH mutant co-deleted, IDH mutant non-codeleted, converges with different patient age at diagnosis distributions, which strongly suggests that the three groups represent distinct gliomagenic biologies.

Gliomas display significant intertumoral and intratumoral heterogeneity: cancer cells from the same tumor cell of origin may contain a wide range of genetic and epigenetic states.^39-41^ Intratumoral heterogeneity confounds diagnosis, challenges the design of effective therapies, and is a determinant of tumor resistance.^42^ Molecular heterogeneity in GBM has been characterized with multiple approaches. For example, fluorescent in-situ hybridization (FISH) analysis of the most commonly amplified receptor tyrosine kinases (RTK) in GBM (*EGFR*, *PDGFRA* and *MET*) revealed a mosaic of tumor subclones marked by different RTK amplifications in 2–3% of GBM,^43-45^ possibly indicating cooperation between cell populations. Single-cell sequencing demonstrated comparable non-overlapping subclonal GBM cell populations marked by different *EGFR* truncation variants, suggesting convergent evolution of *EGFR* mutations.^46^ Genomic profiling of spatially distinct tumor sectors has revealed partial overlap in mutation content in multiple samples from IDH-mutant lower-grade glioma^19,38,47,48^ and IDH-wildtype GBMs.^36,37,49-51^ Mutations/DNA copy number alterations in important glioma driver genes such as *TP53* and *PTEN* have been found to be subclonal, i.e. not present in all cells from the same tumor, suggesting they were acquired after tumor initiation. These unexpected discoveries show the many options tumor cells have to circumvent anti-tumorigenic hurdles such as senescence and geno7c instability. The possibility of extrachromosomal oncogene amplification adds an additional layer of complexity, allowing tumors to rapidly increase intratumoral heterogeneity in response to a microenvironment sparse in resources.^52-57^

Mutation retention rates may be correlated with the geographical distance of samples in the tumor,^51^ and by extension, the level of heterogeneity between different lesions of multifocal GBM is greater than between different areas of the same GBM.^51,58,59^ Spatial heterogeneity determined by genetic alterations is reflected in the epigenetic patterns of different tumor sections examined by combined analysis of DNA methylation and genetic abnormalities.^47,50^ These accumulating data suggest that intratumoral heterogeneity is encoded through a genomic-epigenomic codependent relationship,^47^ in which epigenetic changes may modulate mutational susceptibility in proximal cells, and specific mutations dictate aberrant epigenetic patterns.^47,60,61^ Although gene expression signatures can be used to subclassify GBMs, the predominant subtype often varies from region to region within a given tumor.^37,50^ This relative instability may be in part due to the variable levels of tumor-associated non-neoplastic cells that can be found in different parts of the tumor.^62,63^ Single-cell RNA sequencing of GBM cells confirmed that multiple subtype classifications can be detected in all tumors, often with one subtype dominating the others.^51,63-65^ Transcriptomics and genomics converge at the single-cell level with the observation of mosaic expression of RTK, extending previous observations of mosaic RTK amplification in a small subset of GBM to being a more frequent disease characteristic.^64,65^ Single-cell RNA sequencing further has shown that all gliomas contain cellular hierarchies along an axis of undifferentiated progenitors to more differentiated cell populations, reminiscent of the hematopoietic stem cell hierarchy. The balances shifts towards proliferating progenitors in IDH-wildtype glioma reflecting the clinically more aggressive disease course. ^66-68^ Such developmental and functional hierarchies are associated with dynamic neural stem cell expression patterns in which stem or progenitor cells may function as units of evolutionary selection.^69^

### 2.3 Longitudinal DNA profiling in pre-treatment and post-treatment tumors

One of the earliest reports on the effects of therapy on the tumor genomic landscape analyzed a 23-patient cohort of IDH-mutant lower-grade glioma treated with temozolomide chemotherapy.^70^ A subset of the recurrent tumors acquired hundreds of new mutations that bore a characteristic signature of temozolomide-induced mutagenesis, suggesting that treatment pressure from an alkylating agent induced the growth of tumor cells with new mutations (therapy-induced acquired resistance).^38^ These hypermutated tumors may be sensitive to immune checkpoint inhibitors,^23^ including programmed death-1 (PD-1) inhibitors^71^ and poly-adenosine diphosphate ribose polymerase (PARP) inhibitors (PARPi).^72^ However, clinical trial data supporting these hypotheses have yet to emerge. Another study used whole-genome and multisector exome sequencing of 23 predominantly IDH-wildtype GBM and matched recurrent tumors.^36^ The study showed that some GBM recurrences bore ancestral p53 driver mutations detectable in the primary GBM counterparts (intrinsic resistance), while other recurrences were driven by branched subclonal divergent mutations not present in the parental primary GBM. This may reflect treatment-induced resistance through DNA mutagenesis and a distinct evolutionary process.^36^ As in the study of IDH-mutant lower-grade glioma, a subset of the disease recurrences were characterized by an accumulation of mutations in association with temozolomide treatment. Notably, this effect was limited to cases with *MGMT* promoter methylation. *MGMT* is a gene in the DNA repair pathway, and mutations of other pathway members, such as *MSH2* and *MSH6*, have been nominated as drivers of the hypermutation process.^73^ The spatiotemporal evolutionary trajectory in paired gliomas between initial diagnosis and relapse was further portrayed via integrative genomic and radiologic analyses. Whole-exome sequencing of 38 primary and corresponding recurrent tumors revealed two prevalent patterns of tumor evolution. Linear evolution, in which a recurrent tumor is genetically similar to the initial tumor, was predominantly observed in a subset of recurrent tumors that relapsed adjacent to the primary tumor site. Branched evolution was more common in recurrences at distant sites, which were marked by a substantial genetic divergence in their mutational profile from the initial tumor, with key driver alterations differing in more than 30% of the cases, demonstrating branched evolution. Geographically separated multifocal tumors and/or long-term recurrent tumors were seeded by distinct clones, as predicted by an evolution model defined as multiverse, i.e. driven by multiple subclonal cell populations.^51^

Comprehensive genomic analysis of the processes regulating tumor evolution necessitates serial profiling of pre-treatment and post-treatment tumors. Patients receiving medical care may move to a different medical center in the interval between initial diagnosis and recurrence, which creates significant challenges for the serial collection of tumor tissue. In an effort to elucidate the diverse evolutionary dynamics by which gliomas are initiated and recur, the clonal evolution of GBM under therapy was assessed from an aggregated analysis of datasets generated by multiple institutions.^74^ Systematic review of the exome sequences from 93 patients revealed highly branched evolutionary patterns involving a Darwinian process of clonal replacement in which a subset of clones with selective advantage during a standard treatment regimen renders the tumor susceptible to malignant progression. Mathematical modeling delineated the sequential order of somatic mutational events that constitute GBM genome architecture, identifying mutations in *IDH1, PIK3CA,* and *ATRX* as early events of tumor progression, whereas *PTEN, NF1, and EGFR* alterations were predicted to occur at a relatively later stage of the evolution.^51^ Similar observations have been reported from comprehensive studies of low-grade gliomas, demonstrating that the mutations in *IDH1, TP53,* and *ATRX* were frequently acquired and retained throughout tumor progression from primary to relapse^19,48^. Additionally, these studies have demonstrated crucial insights into oncogenic pathways that drive malignant progression of low-grade gliomas with mutations in *IDH1,* suggesting convergence on aberrantly activated MYC, RB and RTK-RAS- PI3K signaling pathways. Longitudinal profiling of paired samples continues to reveal deeper insights into the genomic background of treatment-induced hypermutagenesis and its potential increased aggressive clinical behavior and relevance in targeted therapy and immunotherapy.^19,48,75,76^ The implications of these data and how these insights can be integrated into clinical practice require further evaluation. Collectively, longitudinal genomic profiling will be essential in implementing clinical application towards patient-tailored treatment regimen.

### 2.4 Transcriptional changes during glioma progression

Unsupervised transcriptome analysis of GBM converged on four expression subtypes, referred to as classical, mesenchymal, neural, and proneural, which are associated with specific genomic abnormalities.^14,15,18,77^ The proneural and mesenchymal subtypes have been most consistently confirmed in the literature, whereas the neural type may simply represent GBM containing a relatively high amount of admixed non-neoplastic neurons.^63,78^. Transcriptional subtyping of the relatively homogeneous IDH-mutant and 1p/19q-codeleted groups have been less emphasized in the literature, as these cases usually carry a proneural signature.^12,14^ While expression subtype classification is a widely used research tool, it has not been shown to correlate with clinical outcome, and has not been incorporated in the recent WHO CNS tumor classification update. Much is still unknown about the drivers of transcriptional subclasses in GBM, their plasticity and how they evolve under therapy. A switch from proneural to mesenchymal expression has been observed upon disease recurrence and proposed as a source of treatment resistance in GBM relapse,^18,79-81^ but the relevance of this phenomenon in glioma progression remains ambiguous, particularly considering a. the increased fraction of microglial/macrophage cells in mesenchymal GBM that confound subtype characterization^62,63^ and b. glioma neurospheres derived from mesenchymal GBM are frequently classified as proneural.^79^ Deriving an expression subtype classification on the basis of glioma-intrinsic genes has maintained the proneural, classical, and mesenchymal classes ^63^. Determining subtypes in a cohort of 91 IDH-wildtype GBM showed subtype switching following therapy and disease relapse in 45% of the cohort ^63^. These patterns converged with changes in the tumor microenvironment, corroborating observations from single-cell transcriptomics that every GBM comprises different subtype mixtures but also revealing that *NF1* loss results in macrophage/microglia recruitment. The ability of genomic abnormalities to regulate the tumor microenvironment suggest cooperation and shows how tumors act as a system or an organ, rather than an aggregation of individual aberrant cells.

### 2.5 Epigenetic changes during glioma progression

DNA methylation profiling of gliomas has prognostic value independent of the age of the patient and the tumors pathologic grade.^11^ Although clonal selection under therapy of genetic mutations nominates mutations as drivers of therapy resistance, strong evidence also suggests that evolutionary selection acting on the epigenome, in the absence of genetic changes, affords plasticity of cells to resist therapy.^11,47^ For example, recurrent IDH-mutant gliomas profiled for mutations and DNA methylation independently evolved deregulation of their cell cycle programs, through genetic mutations or epigenetic mechanisms.^47^ Epigenetic convergence on genetically deregulated biological processes demonstrates that epigenetic abnormalities provide cell subpopulations with fitness advantages that could undermine therapy.

Nearly all IDH-mutant gliomas exhibit a characteristic CpG island hypermethylator phenotype (G-CIMP).^12^ Possible hypotheses for this relationship are as follows (i) DNA hypermethylation induces silencing of key extracellular matrix and cell migration gene promoters,^12^ (ii) DNA methylation mediates alteration of chromosome topography, leading to oncogene upregulation^82,83^ (iii) histone methylation-related changes in gene expression, (iv) DNA hypermethylation associated with mutant IDH may play a role in creating an immunosuppressed microenvironment.^84^

G-CIMP tumors can be further parsed into subsets with reduced genome-wide DNA methylation levels.^11^ While almost all IDH-mutant tumors are G-CIMP at diagnosis, a longitudinal analysis showed that 34% of cases exhibited demethylation towards G-CIMP-intermediate or G-CIMP-low DNA methylation at recurrence.^85^ Substantial epigenetic heterogeneity between tumor samples from the same patient collected at subsequent surgeries was also observed in a cohort of 112 primary GBM patients ^86^. Characteristic trends in DNA methylation between primary and relapsed GBM included a prominent demethylation of Wnt signaling gene promoters, which was associated with worse patient outcome. Moreover, patients whose primary tumors harbored higher levels of DNA methylation erosion showed longer progression-free survival and a trend towards longer overall survival ^86^. This study also explored associations between changes in DNA methylation and magnetic resonance imaging (MRI) and digital pathology data, highlighting the connectedness of the various levels of molecular, cellular and phenotypic heterogeneity in GBM. Analysis of larger cohorts is needed to determine the association between genomic and epigenetic deregulation.

### 2.6 Imaging and (epi)genomics

MRI is a crucial part of standard diagnostic work-up and follow-up of brain tumor patients. It is noninvasive, and owing to the lack of radiation exposure, repeat imaging is not harmful. Conventional MR tumor imaging includes precontrast and post-contrast T1-weighted (T1w) and T2-weighted (T2w)/T2w fluid-attenuated inversion recovery (T2-FLAIR) imaging to assess tumor location, size, and certain macrostructural features.^87^ Newer techniques such as perfusion imaging provide a measure of tumor vascularization in terms of relative cerebral blood volume, which correlates with tumor grade.^88,89^ Relative cerebral blood volume reflects biological behavior of tumors, which might relate to molecular profiles.

In the rapidly growing field of research called radiogenomics,^90^ a rich set of quantitative imaging features are linked with genomic profiles. It has recently been applied in the context of high-grade glioma.^90-92^ Given the major differences in DNA characteristics, gene expression profiles, and DNA methylation profiles, a priority of radiogenomics research on glioblastoma is to identify MR imaging based biomarkers of molecularly defined lower-grade glioma subtypes such as IDH-mutant versus wildtype and 1p/19q codeleted versus non-codeleted. Noninvasive phenotypical assessment has several clear advantages. First, it provides an early test to stratify IDH-mutant, 1p/19q non-codeleted glioma tumors, identifying those patients who are candidates for the most aggressive therapeutical strategies and those for whom a more conservative approach may be preferred. Providing reliable prognostic information through MR imaging can improve patients’ quality of life by postponing, and potentially obviating the need for surgery for a subset of patients.^93^ Second, it would be a means of selecting and tracking patients for personalized treatment regimens in clinical trials.^94^ Third, a detailed global assessment of spatial and longitudinal heterogeneity of gliomas becomes feasible.^95^

## 3. Barriers to progress

The major obstacle to glioma patients currently is a lack of effective treatments, yet we have little understanding of why treatments fail. These failures are likely related to dynamic tumor evolution where treatment-resistant glioma cells are favored over treatment-sensitive cells. As a result, therapy has profound effects on tumor composition by activating intrinsic and adaptive resistance mechanisms, most clearly reflected by the temozolomide induced hypermutator phenotype ^70^. Such processes may be directly induced by the therapy itself, or they are the result of survival and clonal expansion of tumor cells with genetic, epigenetic and/or regulatory alterations that confer drug resistance. Moreover, tumor cells may attenuate the immune response, locally and systemically, to prevent immunological recognition and clearance. All of these processes result in molecular characteristics of the recurrent tumor that differ in significant ways from those found in the primary tumor ^36,38^.

To improve the outcomes of patients with gliomas, we need to explore new therapeutic approaches based on a thorough understanding of treatment-induced molecular and genetic diversity that leads to resistance. The TCGA glioma effort and similar initiatives elsewhere have established comprehensive portraits of the interpatient variability of untreated glioma genomes. Single cell sequencing and barcoding experiments have demonstrated functional hierarchies providing important insights into characteristics of the most relevant cells to target ^66-68^. We are increasingly able to infer the life history of glioma, from tumor-initiating events such as *IDH1* mutation to tumor-promoting events such as RTK alterations. A detailed understanding of the biological diversity within every tumor following clinical presentation and disease progression is needed if we are to successfully understand how treatment affects glioma progression, a needed step towards integration of precision therapeutics into clinical decision making. These considerations also highlight the danger in considering treatment options for patients with recurrent tumors solely on the basis of the molecular analysis of their treatment-naïve tumors. This is particularly important in the setting of clinical research, which often recruits patients with recurrent GBM to evaluate drugs developed on the basis of mechanistic data obtained on treatment-naïve tumors.

Studying heterogeneity and spatiotemporal evolution of cancer in general, and particularly in brain cancer, is challenging. Many tumor samples, and therefore large-scale collaboration, are needed to achieve meaningful comprehensive results. For example, to identify 80% of all somatic alterations occurring in at least 3% of the patient population, a cohort of 500 samples would be needed.^96^ It is crucial to recruit sufficient numbers to validate findings and to capture low-frequency alterations. Individual research groups typically do not have the resources to use a multiplatform analysis of their samples, owing to cost or availability of expertise. Existing longitudinal datasets that have been published consist of a mixture of different modalities, ranging from only exomes^38^ or DNA methylation profiles^47,86^ to a combination of exome sequencing, RNA sequencing, and DNA copy number profiling,^36,63^ thwarting meta-analyses based on cross-publication comparisons. The value of establishing a comprehensive multiplatform reference dataset quickly has been demonstrated by the success of TCGA glioma projects, which have led to a fundamental reclassification of gliomas by the WHO^8^ and are highly cited.^9,12-14^ Similarly, a consortium would be the most effective approach assembling the large cohorts of primary and recurrent pairs needed to identify somatic alterations enriched after disease progression. Systematizing and standardizing what we do and how we do it will be essential for affecting paradigmatic change to clinical practice in neuro-oncology. This philosophy is at the core of the international GLASS consortium.

## 4. The Glioma Longitudinal AnalySiS (GLASS) consortium

The rapidly growing body of knowledge in molecular and genomic data is refining clinical diagnosis and prognostication (exemplified by the revised fourth edition of the WHO classification of CNS tumors published in 2016^8^), but has not resulted in improvements clinical outcomes, particularly in the more aggressive gliomas. This is evidenced by numerous trials that have failed to reach their primary endpoint.^97,3,98^ These successes and failures highlight the need for large-scale collaborations that to help us understand the impact of treatment on evolutionary dynamics and, most importantly, why treatments fails?

The recognition of the need is why we initiated the GLASS consortium: to achieve power from the use of large numbers of paired samples from all contributors. GLASS aims to perform comprehensive molecular profiling of matched primary and recurrent glioma specimens from an unprecedented 1500 patients, 500 in each of the three major glioma molecular subtypes: 1. IDH-wildtype; 2. IDH-mutant; and 3. IDHmutant with 1p/19q codeletion. The consortium at the time of writing includes investigators from 32 academic hospitals, universities, and research institutes from 12 countries, (see list of participants on the GLASS website, http://www.glassconsortium.org). By analogy with the International Cancer Genomics Consortium ^99^ GLASS is structured into country-specific franchises (GLASS-NL, GLASS-AT, GLASS-AU, GLASS-Korea, etc.) led by local investigators who are invested in the team’s overall goal of assembling a meaningful sample cohort of pretreatment and post-treatment samples for each glioma grade and type, while taking advantage of country-specific opportunities. This enables each GLASS branch to have unique features that allow deeper analysis of subcohorts, that is, with additional imaging annotation, parallel characterization of drug response through xenografting of tumor samples, specific focus on a glioma subtype, etc., thereby making them competitive and enabling them to address non-overlapping aspects of the phenotypic diversity seen in the clinic. Country-specific branches will be coordinated to connect with the larger analyses, to drive specific research topics for both. There are no explicit restrictions on publishing, that is, each group is invited to publish their substudies independently. The overall goal is to establish a reference data set by pooling samples and aggregate data from all multiplatform analyses, countries and substudies, and to make datasets comparable through coordinated sample and data processing guidelines. Country franchises are centrally connected through a number of committees, each overseeing different aspects of the analysis (Figure 2).

**Figure 2.**
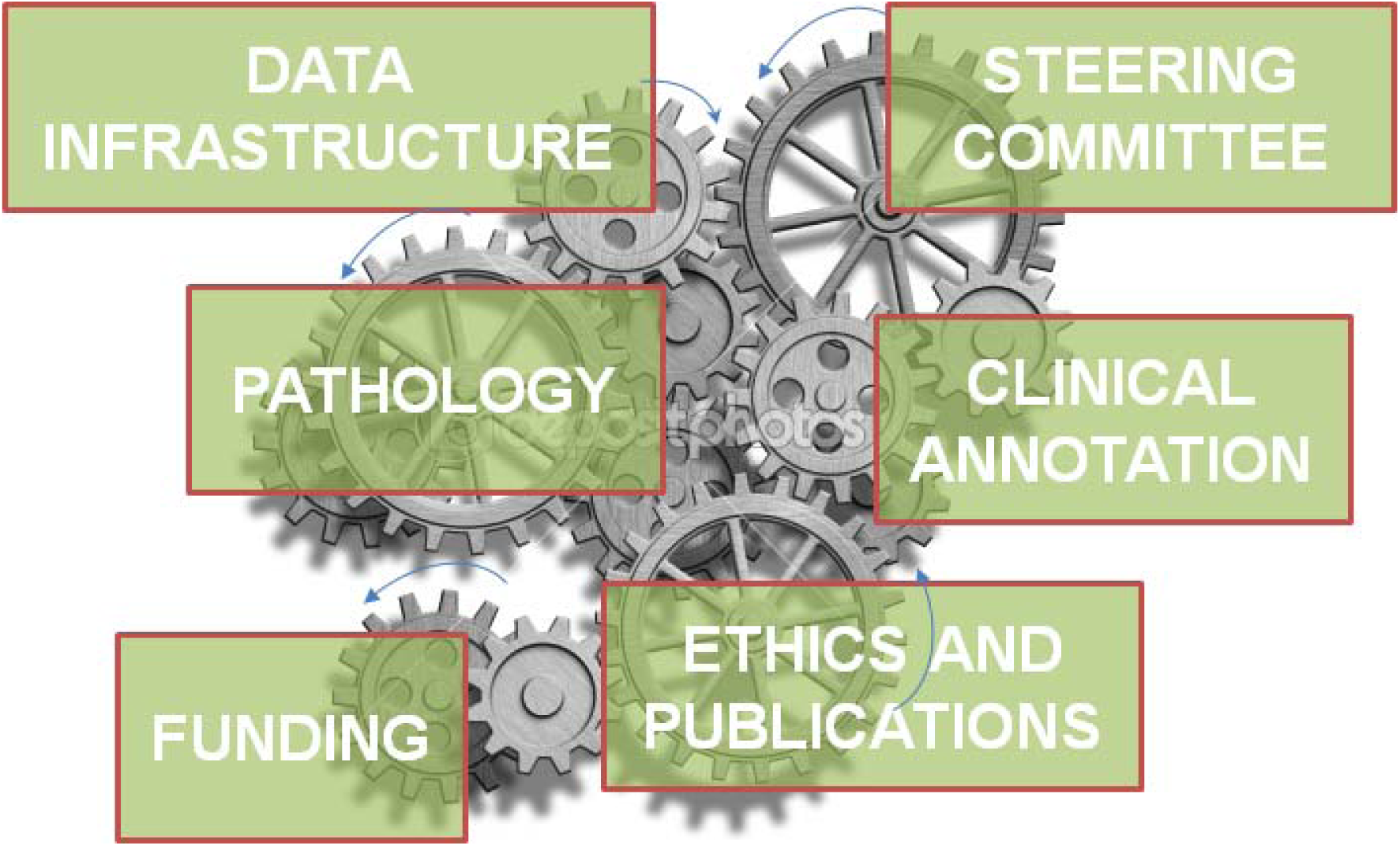
Overview of GLASS committees. Details on committee mandates are provided in section 4.

### 4.1 Biospecimens and characterization platforms

Biospecimens from gliomas are often snap frozen or conserved formalin fixed, paraffin embedded (FFPE). For genomic and transcriptomic analyses, snap frozen material is preferred, while historically FFPE is the common approach to tissue preservation. Methods for generating sequencing data from FFPE material are increasingly improving, with 5–20% of samples failing quality controls. Given that samples from multiple timepoints are required for inclusion into GLASS, patients for whom only FFPE material is available are twice as likely to not yield sufficient high quality DNA. RNA extracted from glioma tissue is often highly degraded resulting in higher attrition rates^100^, but high quality RNA sequencing data from FFPE samples has been reported ^101^. For DNA methylation profiling of FFPE material, a recent study focusing on primary glioblastoma reported a high success rate using the reduced representation bisulfite sequencing assay ^86^. While we require the availability of a matching germline sample (often but not always from blood) for inclusion of DNA sequencing data into GLASS, cases without a germline match may be candidates for transcriptome and DNA methylation analysis. Ideally, we aim to generate DNA, RNA and epigenomic sequencing data from every patient. Single-cell analysis methods require fresh tissue from which individual cells can be dissociated; they are currently outside the scope of GLASS, but may be considered in the future as the project evolves or as part of specific subprojects. Similarly subsets of the GLASS cohort will be compared longitudinally by spatial correlation using multisector analysis (3–6 samples per tumor) to understand whether any differences between paired tumor samples are the result of intratumoral heterogeneity or longitudinal heterogeneity. Where available, these will be correlated with conventional and novel MR imaging to explore spatiotemporal heterogeneity noninvasively. We aim to take current radiogenomic approaches further, not only to establish features of genetic characteristics at first diagnosis, but also in relation to molecular alterations over time (including under pressure of standard therapy).

Comprehensive genomic sequencing is needed to identify patterns of disease evolution as well as key mutations and chromosomal alterations that confer resistance to standard radiation, temozolomide, and novel clinical trial therapies. Sequencing paradigms and their costs are rapidly evolving, and each method provides different but often complementary information. There is no consensus on optimal methods. With the accessibility of $1000 per biospecimen whole genome sequencing (WGS), the costs of WGS and whole exome sequencing (WES) have become comparable. WES has better sensitivity in detecting mutations in coding regions, but does not interrogate noncoding regions of the genome, structural variants, or noncoding copy number variants. The comprehensive nature of WGS enables analysis of evolution and clonality at higher resolution. WGS and WES combined may provide the optimal window on the breadth, depth, and allelic fraction of somatic events. However, where limitations in tissue or resources mandate a choice of one or the other, the decision will depend on the purpose of the (sub) project.

### 4.2 GLASS committees

GLASS has established different committees with the expertise to coordinate the various aspects of the consortium. They include pathology, clinical annotation, data infrastructure, ethics and publication, and funding committees.

#### 4.2.1 The GLASS pathology committee

maintains centralized classification and tissue processing. The committee has set up a panel of specific inclusion criteria for tissue samples. A prerequisite for inclusion is that the patient has given informed consent to donate tissue to research and that an adequate blood sample or any other adequate source of genomic DNA is available. Both FFPE and snap-frozen tissue samples will be included in the GLASS study and evaluated by the committee. The area of viable tumor tissue should be at least 50 mm^2^ and the tumor cell percentage should be higher than 50%. Hematoxylin and eosin slides of each sample will be digitized by an automated slide scanner and the images will be stored on a central server in order to be accessible by all members of the GLASS expert pathology committee. An anonymized pathology report of both the original and the recurrent tumor has to be submitted for review to the committee with information on microscopic (including immunohistochemical) findings and, if performed, results of molecular analyses, as well as the integrated diagnosis. On the basis of this information, the GLASS expert pathology committee will formulate a (tentative) review diagnosis and thereby select patient samples that can be used for further study by the GLASS consortium. Corresponding whole-slide images of all patient samples that are included in GLASS will then be made available as a digital resource for further image analysis ^102,103^.

#### 4.2.2 The GLASS clinical annotation committee

maintains standardized data processing, data management and data sharing. The currently available large-scale datasets suffer from relatively weak clinical annotation. Consequently, linkage of genotype with clinical and morphological phenotype remains to be fully exploited in primary and recurrent settings. The GLASS clinical annotation committee will address this by standardizing clinical and imaging data collection for prospective studies and oversee aggregation of the clinical and imaging data from patients whose profiles are already included in the composite dataset. Insight into mechanisms of response and resistance and exposed therapeutic vulnerabilities will be fed into current and future clinical trial designs by GLASS investigators and trials designed collaboratively in academia and industry or across both. This strategy will require an integrated bioinformatics interface across the molecular and clinical research. The necessary data processing infrastructure will be developed by the GLASS consortium and distributed among its franchises, to ensure compatibility, comparability, and reproducibility. Each individual franchise will make clinical, imaging, and molecular data accessible in a comprehensible way by integrating clinical, imaging, and molecular parameters to explore correlation with relapse data. Currently, radiology and imaging are part of the clinical annotation committee. By mapping imaging features in a voxel-wise manner and correlating these spatially with molecular alterations obtained from different parts of the tumor we aim to assess the entire tumor and to determine intratumoral heterogeneity.

#### 4.2.3 The GLASS data infrastructure committee

maintains standardized data processing, data management and data sharing. A characteristic of the GLASS consortium is that data will be generated at multiple institutions distributed over multiple countries. As the regulations pertaining to ethical use of sequencing datasets are continuously evolving, GLASS will follow the example set by ICGC to perform decentralized data analysis to avoid cross-border exchange of patient-sensitive raw sequencing data. The GLASS data infrastructure committee has developed Docker software images that can be shared by participating institutions to ensure analysis uniformity. Like a shipping container, a Docker image packages one or more software tools to establish a workflow resembling an executable application. Comparable to platform-independent Java software, the ready-to-run Docker images are independent of the local computational environment. The GLASS participants run the Docker image locally, which initializes a per-sample-per-analysis Docker container, resulting in data analysis using an identical software environment and run parameters. Docker images are available to process exome sequencing data, which includes alignment, quality control, mutation calling, and DNA copy number estimation. Comparable Docker images are ready for processing of whole-genome sequencing and transcriptome sequencing data. Docker images are available for download through http://docker.glassconsortium.org *(RV: currently pending)*.

The data infrastructure committee will also coordinate mechanisms for dissemination of results, as to widely share datasets with the community. We may explore mechanisms such as the Genomic Data Commons, or similar, in order to align our efforts with other molecular profiling studies.

#### 4.2.4 The GLASS ethics and publications committee

was created to identify and address critical ethical, legal and social questions faced by researchers and patients participating in the GLASS program. The guidelines established by this group will continue to inform future policies that ensure effective and fair use of cancer genomic information coupled with relevant clinical annotations. All participating sites in the program have institutional ethics approval for data protection and the use of tissue and/or DNA and clinical and where applicable imaging data from patients that have given written informed consent. Data will be made available rapidly after generation for community research use. Publication guidelines will follow that established by the TCGA policy for authorship and publication.

## 5. Final remarks and perspectives

Survival and quality of life for patients with diffuse gliomas remains dismal with standard treatments. Diffuse glioma is a fatal disease with an enormous societal burden as a result of short survival following high-grade disease and the relatively young age at diagnosis of lower-grade disease. This not only affects patients in the prime of their life, but also puts enormous burden on their immediate entourage, as they need extensive supportive care and navigation through a complicated medical landscape, and difficulties with medical costs and insurance. While cures of diffuse gliomas remain elusive, our patients demand better therapies and, with no substantive impact of molecular medicine, in practice treatments remain a ‘one size fits all’. The GLASS Consortium will enable the improvement of clinical outcomes by establishing a broadly useful reference data set that will provide pivotal new insights into mechanisms used by gliomas to defy therapeutic challenges. For example, hypermutation following temozolomide treatment occurs in up to 15% of glioma, but too few samples have been profiled to understand what is driving this process or to identify biomarkers predictive of a TMZ associated hypermutator phenotype. GLASS will have the power to identify molecular markers indicating evolution from newly diagnosed to highly aggressive therapy-resistant malignancy, in addition to molecular aberrations that occur under pressure from different therapeutic modalities. It will also allow for the identification of currently undiscovered molecular targets for resistance-prevention agents that might be co-administered with classical therapies. Systematic correlation of the molecular information with clinical, imaging, and pathology data will help improve interpretation of prognostic findings in the course of the disease.

Finally, and importantly, GLASS is an opportunity for interchange of knowledge among an international group of collaborators to ultimately build smarter clinical trials and develop therapies that will extend survival and improve the quality of life of people with diffuse gliomas. GLASS is well positioned to demonstrate the value of well-coordinated collaborative efforts. To that extent, new investigators are invited to join to the Consortium, where major criteria for participation are the ability to offer datasets of longitudinally profiled glioma patients or the availability of suitable tissue samples.

In summary, we hope that through the GLASS Consortium, we continue the immeasurable success of The Cancer Genome Atlas while increasing the focus on making a difference to patients and their families. Therapy resistance is what kills patients, and GLASS will inform on how to avoid it.

## Acknowledgements

The GLASS Consortium is indebted to the support from the National Brain Tumor Society, Oligo Research Fund, and Alan and Ashley Dabbiere (RGWV, JC). Rhana Pike at the National Healh and Medical Research Council Clinical Trials Centre, The University of Sydney, Australia provided editorial assistance. GLASS-AT is supported in part by Austrian Science Fund grant KLI394 (AW), by the Austrian Academy of Science (CB), and by the European Research Council (European Union’s Horizon 2020 research and innovation program, grant agreement no. 679146) (CB). GLASS-L is supported by a Télévie grant (SPN). GLASS-NL receives funding from the Dutch Cancer Society KWF, grant 11026 (PW). GLASS-USA has been supported by the Cancer Prevention and Research Institute Texas, grant R140606 (RGWV, JH, JDG). Tissue procurement at the Winship Cancer Institute is supported by NIH-NCI P30-CA138292 (EVM). Support is also provided by a Henry Ford Hospital institutional grant, grant 2015/07925-5, 2016/15485-8, 2014/08321-3 from Sao Paulo Research Foundation (FAPESP)(HN). The Else Kröner-Fresenius-Stiftung (2015_Kolleg_14) supports clinical and tissue database at the University Tübingen (GT).

